# Mitochondrial Dysfunction and Senescence Accompany Glioblastoma Cell Death Triggered by a Putative Metabolic Inhibitor

**DOI:** 10.1101/2025.09.08.674975

**Authors:** Anjali Yadav, Shraddha Bhutkar, Shrikant Barot, Vikas Dukhande

## Abstract

Glioblastoma (GBM) is a fatal cancer with a dismal prognosis and a dire need for novel chemotherapeutics. Metabolic reprogramming is an established hallmark of cancer. In our previous study on GBM, we aimed at targeting the metabolic reprogramming of cancer by using stiripentol (STP), a putative lactate dehydrogenase (LDH) inhibitor and an FDA-approved anti-epileptic drug. However, the precise mechanism of STP’s anti-cancer activity remains unclear. We aimed to elucidate the mechanism of action of STP in GBM to further develop STP as a therapeutic. We employed a multiomic approach followed by metabolic and cellular assays. STP treatment induced genetic and metabolic alterations in GBM cells. Inhibition of LDH by STP was moderate but not potent. The cellular changes were accompanied by an increase in reactive oxygen species, a decrease in mitochondrial membrane potential, and induction of senescence in GBM cells. Our research indicates that further research in senescence-inducing agents and novel LDH inhibitors can provide novel therapeutics for GBM.

## Introduction

Glioblastoma multiforme (GBM) is one of the most aggressive primary malignant brain tumors. It accounts for approximately half of all primary central nervous system tumors [1]. The 5-year survival rate of GBM is ∼7% and the median survival time of patients diagnosed with GBM is merely 12 to 15 months [2]. GBM is characterized by rapid proliferation, a highly invasive nature, tumor heterogeneity, and resistance to chemotherapy. The traditional treatment modalities for GBM include surgical resection, chemotherapy, and radiation therapy [3]. Despite the advancements in the search for new therapeutics, there has not been any breakthrough. The presence of the blood-brain barrier (BBB) coupled with heterogeneous composition, rapid turnover, epigenetic alterations, and metabolic reprogramming contributes to the limitation in development of new therapeutic [4].

One of the major hallmarks of cancer is the tumor’s ability to rapidly reprogram its cellular metabolism to support growth and survival [5]. GBM cells meet their increased energy and biosynthetic demands by metabolic reprogramming. Under normoxia, pyruvate would enter the TCA cycle via acetyl-CoA, generating NADH and FADH_2_ for oxidative phosphorylation, which would yield ATP for energy consumption. Under hypoxia, anaerobic glycolysis takes over, converting pyruvate to lactate and yielding less ATP. When cells exhibit the Warburg effect, they adapt to rapid ATP generation via glycolysis, even in the presence of oxygen, also termed aerobic glycolysis [6]. The shift to aerobic glycolysis yields lactate as a byproduct. The process of aerobic glycolysis allows the biosynthesis of the building blocks for cellular proliferation [7]. It also creates an acidic tumor microenvironment that allows the survival of GBM cells. One of the key metabolic reprogramming enzymes involved in aerobic glycolysis is lactate dehydrogenase (LDH). LDH interconverts pyruvate to lactate, regenerating NAD^+^. There are several studies emphasizing the inverse correlation between LDH expression and patient survival [8]. Specifically, elevated LDHA activity has been linked to tumor progression and resistance to several chemotherapies [9]. This makes LDHA inhibition a promising target for a novel therapeutic for the treatment of GBM.

In our previous study, we explored the potential of multiple metabolic inhibitors for their effect on GBM. Our research led us to stiripentol (STP), an FDA-approved anti-seizure drug used for Dravet’s syndrome-a rare kind of epilepsy [10]. Although the putative mechanism of STP’s anti-seizure activity is GABA-A potentiation, it has been recently studied for its potential as an LDHA inhibitor [10–12]. We studied STP in GBM cells and discovered its *in vitro* efficacy [10]. However, the mechanism of STP’s anti-cancer activity is not known. In this study, we aimed to decipher the mechanism of STP’s anti-cancer activity in GBM using a multiomic approach encompassing transcriptomics and metabolomics. In addition, we probed metabolic alterations and the cell death mechanisms induced by STP in GBM cells. Our findings revealed some critical insights into the metabolic reprogramming of GBM and the potential of repurposing STP as a therapeutic intervention.

## Materials and methods

U87 cell line was purchased from ATCC (USA), fetal bovine serum (FBS) (Atlanta Bio, USA), Dulbecco’s modified Eagle medium (DMEM), PSA, and Trypsin (Corning, USA), stiripentol and temozolomide (Fisher, USA), GSK2837807 sodium oxamate (Sigma, USA), siLDHA and Lipofectamine RNAimax (Life technologies, USA), LDHA plasmid (Origene, USA), Transit LT-1 (Mirrus, USA), Opti-MEM (Gibco, USA), BCA (Thermoscientific, USA), Direct-Zol RNA extraction kit (Zymo research, USA), CMH_2_DCFDA and CMXROS (Invitrogen, USA), primary antibodies β-actin, ALDHA13, AOX1, CPA4, TXNIP, KYNU, (Proteintech, USA), LC3 (Sigma) secondary antibodies anti-mouse, anti-rabbit, and senescence kit (Cell signaling, USA).

### Cell Culture

The human cell line U87 was obtained from ATCC (American Type Culture Collection). The cells were maintained in DMEM 1 g glucose media supplemented with 10% FBS and 1% PSA at 37 °C with 5 % CO_2_ incubation.

### Overexpression of LDHA

1 × 10^6^ U87 cells were plated in a 100 mm dish with DMEM 1 g glucose media supplemented with 10% FBS and 1% PSA. The cells were incubated overnight in 5% CO_2_ and 37 °C. The following day, LDHA DNA was transfected into the cells. Transit LT1 was used as a transfection reagent with 5 μg and 10 μg of DNA in Opti-MEM no serum media. A GFP control group was added to ensure the transfection efficiency. After 24 h of transfection, lysates were prepared, and protein expression was measured using immunoblotting.

### Knockdown of LDHA

1 × 10^5^ cells were plated in a 6-well plate with 2 mL fresh media. The plate was incubated overnight in 5% CO_2_ and 37 °C. The following day LDHA siRNA was added with the final concentration of 125 nM/well and 200 nM/well. Lipofectamine RNAimax was used as a transfection reagent for the knockdown. Serum-free Opti-MEM was used for the experiment. After 48 h of transfection, the cells were lysed with modified RIPA buffer and 4X SDS dye was added. The lysates were probed for protein expression analysis.

### Western blot

Western blotting was performed to detect proteins in U87 cells treated with STP, TMZ, and their combination. 1 × 10^6^ cells were plated in a 10 cm dish and incubated overnight at 5% CO_2_ and 37 °C. The next day, cells were treated, and protein samples were extracted from the cells using modified RIPA (radioimmunoprecipitation assay) lysis buffer, and the protein concentration was determined using BCA assay. The protein samples were then mixed with a 4X loading buffer and heated to denature the proteins. Equal amounts of protein were loaded onto an SDS-PAGE gel and subjected to electrophoresis. Proteins were then transferred from the gel onto a polyvinylidene fluoride (PVDF) membrane. The membrane was blocked with 5% non-fat dry milk to prevent non-specific binding.

After blocking, the membrane was incubated with a primary antibody overnight. Following washing to remove excess antibody, the membrane was treated with a secondary conjugated antibody. The presence of the target protein was detected using a chemiluminescent substrate using Azure Imaging system (CA, USA). The expression levels were quantified using Image J and plotted on GraphPad Prism software.

### Cell viability

5000 cells/well were plated in a 96-well plate. The cells were incubated overnight in an incubator with 5% CO_2_ and 37 °C. The following day, the cells were overexpressed or knocked down with LDHA for 24 and 48 h. Thereafter, the cells were treated with STP 10, 20, 40, 60 μM along with a positive control, LDH inhibitor GSK2837807A 80 μM. After 48 h of treatment, MTT dye was added, and the cells were incubated with the dye for 3 h. After 3 h of incubation, the formazan crystals were dissolved in DMSO, and the OD value was recorded in a spectrophotometer at 570 nm. The results were quantified and plotted on GraphPad Prism software.

### Enzyme activity assay

1 × 10^6^ U87 cells were plated in a 10 cm dish. The cells were incubated overnight in 5% CO_2_ and 37° C. The next day, cells were treated with STP at 25-, 50-, 100-, 250-and 500-μM concentration and positive control sodium oxamate 500 μM. The cells were then lysed with HEPES Tris buffer. Master mix was prepared with HEPES Tris, NADH, Triton X-100, and H_2_O. The lysates were then added to a 24-well plate. LDH was then added per well, and the uninitiated reaction was measured for 10 min with readings every min. Next, Na pyruvate was added as an initiator, and the kinetic reading for the initiated reaction was measured. The enzyme activity was normalized with the protein concentration. The quantification was then normalized with the protein concentration.

### Metabolomics

1 × 10^6^ cells were plated in a 100 mm dish in DMEM 1 g glucose media with 10% FBS and 1% PSA. The cells were treated with STP 50 μM, TMZ 100 μM, and a combination of STP and TMZ at 50 and 100 μM, respectively for 24 h. Next, the cells were washed

with 100 mM ice-cold ammonium acetate. Cells were then lysed with 100% ice-cold methanol and scraped. The samples were added with internal standards and then frozen under liquid nitrogen and thawed in an ice-cold sonication for 3 cycles. The samples were then centrifuged at 14000 rpm for 10 min, the supernatant was dried under gentle nitrogen flow and reconstituted into 100 μL of 80% methanol for iHILIC column analysis. Next, the samples were dried and reconstituted into 100 μL of 20% methanol for PFP column analysis for widely targeted small polar metabolite panel using Sciex 6500+ QTRAP at Mount Sinai Stable Isotope and Metabolomics Core facility. The raw results were received and analyzed using MetaboAnalyst 6.0 software. Statistical analysis (one factor), enrichment analysis and pathway analysis, and network analysis were performed.

### Seahorse metabolic assay

5000 cells/well were plated in 6 wells of an 8-well Seahorse plate, leaving top and bottom well empty. DMEM 1 g glucose media supplemented with 10% FBS and 1% PSA was used to plate the cells. The cells were incubated overnight in an incubator with 5% CO_2_ and 37 °C. The following day, cells were treated with STP at 25 μM and 50 μM for 24 h. The media was changed with Seahorse Glycostress test and Mitostress test media with several washes as per manufacturer’s protocol [13,14]. Cells were then incubated in no CO_2_ incubator for 20 min. The cartridge was hydrated with a calibrant overnight before the experiment. On the day of assay, after 24 h of drug treatment, Glucose 5.5 mM, Oligomycin 3 μM, and 2-deoxy glucose 50 mM were added to the ports in the cartridge for glycolytic stress test. Oligomycin 3 μM, FCCP 2.5 μM, and rotenone/antimycin A 50 mM were added to the injection ports for Mito stress test. The assays were run on Agilent Seahorse XF mini analyzer. The results were then normalized by performing a protein concentration analysis through a BCA assay. The graph was plotted on GraphPad Prism 9 software.

### Transcriptomics

1 × 10^6^ U87 cells were plated in a 100 mm dish in DMEM 1 g glucose media with 10% FBS and 1% PSA. The cells were treated with STP 50 μM, TMZ 100 μM, and a combination of STP and TMZ at 50 and 100 μM, respectively, for 24 h. RNA was extracted with DNA digestion using the Zymo Research RNA extraction kit. Total RNA quality and yield were determined by the A260/A280 and A260/A230 ratios by nanodrop. The samples were further analyzed at Yale Center for Genome Analysis, where RNA integrity was determined by an Agilent Bioanalyzer or fragment analyzer gel, and the ratio of ribosomal peaks was measured. Samples with RIN values of 7 or greater were used for library prep. For RNA seq library preparation, mRNA was purified from normalized inputs between 10-1000 ng of total RNA using mRNA capture beads prior to fragmentation (Watchmaker mRNA library prep kit). Following first-strand synthesis with random primers, second-strand synthesis and A-ligation with 3’ T overhangs were ligated to library insert fragments. Library amplification amplified the fragments carrying the appropriate adaptor sequences at both ends. Strands marked with dUTP were not amplified. Indexed libraries that met the appropriate cut-off for both were quantified by qRT-PCR using a commercially available kit (KAPA Biosystems) and insert size distribution was determined with the LabChip GX or Agilent TapeStation. Samples with a yield of >0.5 ng/μl were used for sequencing. Sample concentrations were then normalized to 2.0 nM and loaded onto Illumina NovaSeq X plus flow cell at a concentration that yields 20 million passing filter clusters per sample. Samples were sequenced using 150 bp paired-end sequencing on Illumina NovaSeq according to Illumina protocols. The 10 bp unique dual index was read during the additional sequencing reads that automatically follow the completion of read 1. Data generated during the runs were simultaneously transferred to the YCGA high-performance computing cluster. A positive control provided by Illumina was spiked into every lane at a concentration of 0.3% to monitor sequencing quality in real time. Raw FASTQ. GZ files were received, and quality control was ensured using Brahmamatic software. EdgeR software was used to analyze the differentially expressed genes using the RNA pipeline, including Trimmomatic to process and remove adapter sequences of low-quality reads. The cleaned reads were aligned with the reference genome GRCh38. Differential gene analysis was conducted using R DESeq2 to identify differentially expressed genes across all the treatments. Volcano plots, heatmaps, GSEA gene ontology, and pathway enrichment were performed for biological processes and molecular functions associated with the differentially expressed genes. MetaScape analysis was also used to explore pathways involved with significantly altered genes amongst all treatment groups. Adjusted P value <0.05 was used as a strict cut-off for all the above-mentioned analyses. Log fold 1 or above in addition to adjusted p value <0.05 was used to filter the leads, as well as to further validate them.

### ROS assay

2 × 10^4^ cells were treated with drugs for 24 h. Thereafter, the cells were stained with CMH_2_DCFDA dye as per manufacturer’s protocol. Cells were also stained with DAPI for the staining of the nuclei. Cells were incubated for 30-45 min at 37 °C and 5% CO_2._ Next, fluorescent DAPI, GFP, and merge images of the cells were taken on the EVOS FL Auto 2 at 100× magnification. The cells that took up the dye were counted on Image J Fiji software. The graphical quantification was performed on GraphPad Prism software.

### Mitochondrial membrane potential assay

5 × 10^3^ cells were treated with drugs for 24 h. Thereafter, the cells were stained with CMXROS dye as per the manufacturer’s protocol. Cells were stained with DAPI for nuclear staining. Cells were incubated for 30-45 min at 37 °C and 5% CO_2._ Following incubation, fluorescent DAPI, RFP, and merge images of the cells were taken on the EVOS FL Auto at 100× magnification. The cells that took up the dye were counted on Image J Fiji software. The graphical quantification was performed on GraphPad Prism software.

### Lipid peroxidation assay

Approximately 1 × 10^6^ U87 cells were plated in a 10 cm dish in 1 g glucose DMEM media, supplemented with 10% FBS and 1% PSA, and incubated at 37 °C and 5% CO_2_ overnight. The following day, the plates were treated with STP 50 µM, TMZ 100 µM, a combination for 24 hr, and a positive control, H_2_O_2_ at 500 µM for 4 hr. After completion of the treatment, lysates were prepared as per the manufacturer’s protocol. The lysates were normalized using BCA assay. MDA detection reagent was added to the lysates and samples were heated at 95 °C for 30 mins, centrifuged at 4 °C at 13000 rpm for 10 min. Supernatant was added to a 96-well plate, and absorbance was measured using SpectraMax M5e and analyzed on GraphPad Prism.

### Phospho-kinase array

Approximately 1 × 10^6^ U87 cells were plated in a 10 cm dish in 1 g glucose DMEM media, supplemented with 10% FBS and 1% PSA. The plates were incubated at 37 °C and 5% CO_2_ overnight. The following day the plates were treated with STP 50 mM for 24 hr. After completion of the treatment, lysates were prepared as per the manufacturer’s protocol. The lysates were normalized using BCA assay. The lysates were then probed on the nitrocellulose membranes provided by the manufacturer using the Proteome Profiler Human Phospho-Kinase Array Kit (R&D systems). The blot was incubated at 4 °C overnight. The day after, the blots were given washes with the wash buffer provided, and the secondary antibody was added to the blot. The blot was incubated for 1 h at room temperature. Chemiluminescent agents were added, and the blots were read on the EVOS FL Auto 1. Image analysis was performed on Image J software and quantified using GraphPad Prism software.

### Senescence assay

10 × 10^3^ cells were incubated with drugs for 48 h. After the completion of the treatment, the cells were fixed and stained with β-galactosidase staining reagent dye as per the manufacturer’s protocol. Cells were incubated without CO_2._ The images of the cells were taken on the EVOS FL Auto 1 at 200× magnification. The cells that took up the stain were counted on Image J Fiji software. The graph was quantified on GraphPad Prism software.

### Statistical Analysis

All experiments were performed at least in n=3, and statistical analysis of one-way ANOVA Dunnett and two-way ANOVA Tukey test were performed as required on GraphPad Prism software *p <0.033, **p< 0.002, ***p <0.001. All values are expressed as mean ± SEM as mentioned.

## Results

### STP exhibits LDHA inhibitory effects comparable to an established inhibitor

To confirm whether the effect of STP on GBM cells is LDHA-dependent, the cells were overexpressed and knocked down for LDHA using a plasmid and siRNA. Overexpression (OE) of LDHA using a plasmid resulted in approximately a 100% increase in LDHA protein expression in U87 cells (Fig. 1A). However, the transfection by itself decreased the viability of U87 cells (Fig. 1B). The cells with modulated LDHA profile were then treated with STP at different concentrations or with positive control LDH inhibitor GSK 2837808A, and TMZ, and processed for a cellular viability assay. The LDHA OE cells showed no significant difference in cell viability compared to the OE control. Whereas the cells with endogenous LDHA showed significantly decreased cell viability of U87 cells after treatment with STP at 10, 20, 40, and 60 μM, GSK28837808A 80 μM, and TMZ 200 μM for 48 h (Fig. 1B). The data indicates that LDH expression may play a role in STP’s effect on cell viability in U87 cells. This could indicate either an insufficiency of STP to inhibit the significantly higher amount of available LDHA or a compensatory mechanism of U87 cells using an alternative pathway to promote survival. Vice versa, knockdown of LDHA performed by siLDHA showed significantly decreased protein expression of LDHA by 90 to 100% at 200 nM concentration, as shown in Fig. 1C. The silencing of LDHA showed a significant decrease in the viability of U87 cells in comparison to LDHA endogenous cells (Fig. 1D). The cell viability assay performed on LDHA knockdown cells demonstrated an insignificant reduction in cell viability in comparison to LDHA knockdown control, except for TMZ treatment (Fig. 1D). The transfection and silencing effects were strong without treatment, which may confound the cell viability comparisons. After the overexpression and knockdown of LDHA suggested at least a partial role of LDH in STP’s action, we performed an LDH enzyme activity assay at STP concentrations of 25 μM, 50 μM, 100 μM, 250 μM, and 500 μM. A Positive control, sodium oxamate 500 μM, was used. The effect of STP on LDH activity in U87 cells was moderate and significant LDH inhibition was only observed at higher concentrations of STP at 250 and 500 μM (Fig. 1E). Although STP’s anti-seizure activity is putatively attributed to GABA-A receptor (GABRA1) potentiation, the role of GABRA1 in STP’s anticancer activity is unclear. To investigate the effect of STP on GABRA1 as a target for its anti-cancer activity, we performed a cell viability assay in SK-N-SH neuroblastoma cells and used diazepam as a positive control that inhibits GABAergic neurotransmission. Our results showed that STP does not affect the cellular viability in SK-N-SH cells (Fig. 1F).

**Figure 1.**
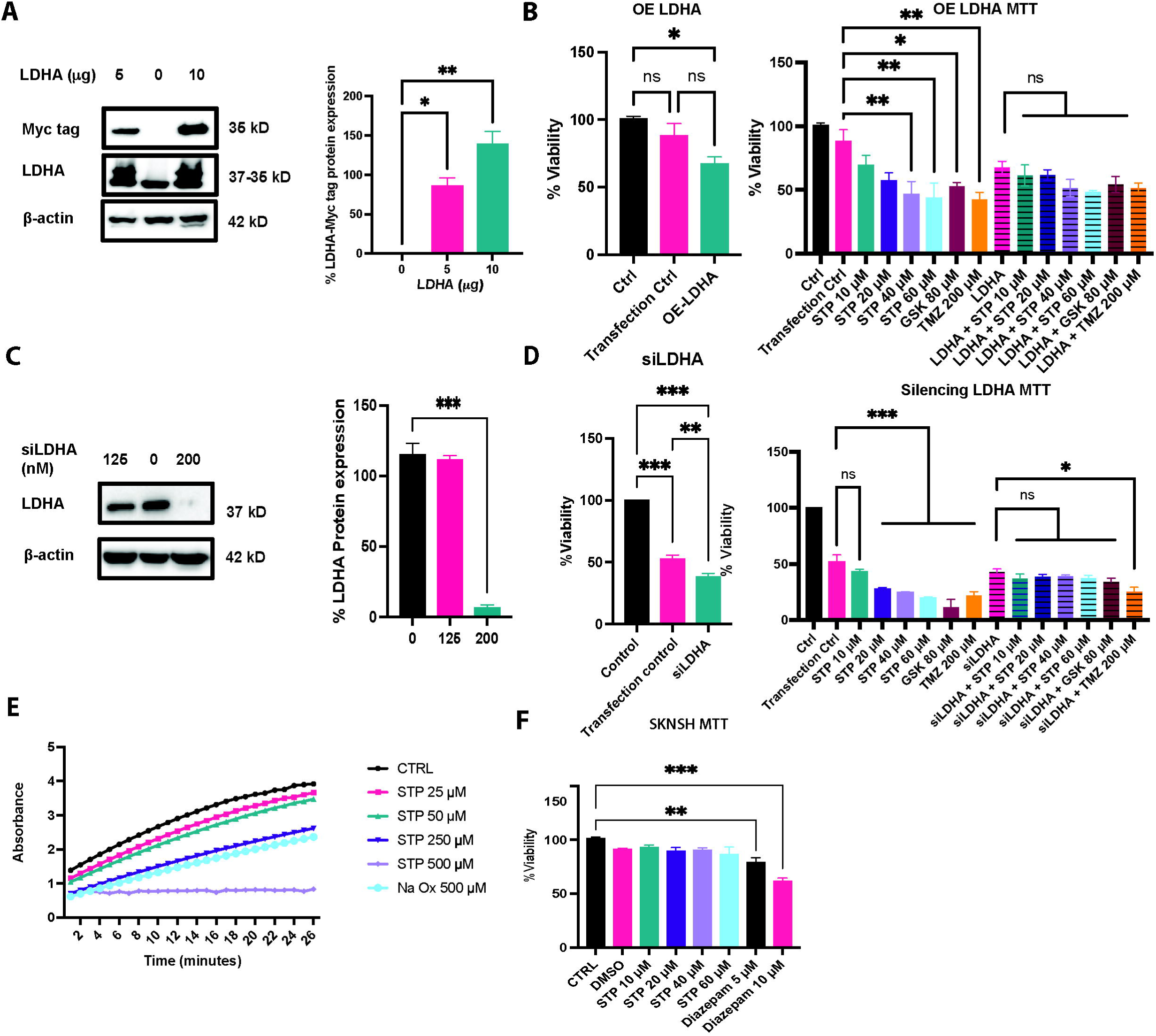
Effect of STP on putative targets LDHA and GABRA1. (A) Immunoblot showing overexpression of LDHA at 5 and 10 mg concentration of the LDHA plasmid in U87 cells, represented with LDHA and Myc tag LDHA. Quantification of the percentage of overexpression observed. (B) Cell viability of control, empty vector, and LDHA plasmid overexpression. Cell viability graphs of STP, TMZ, and GSK without overexpression of LDHA and with overexpression of LDHA. (C) Immunoblot representing LDHA knockdown by siLDHA. Quantification of the percentage decrease in LDHA expression. (D) Cell viability analysis of STP, TMZ, and GSK upon knockdown of LDHA. (E) Kinetic evaluation of LDHA enzyme activity assay with STP, and sodium oxamate. (F) Cell viability of SKNSH cells upon treatment with STP and Diazepam. n=3, *p<0.033, **p<0.002, ***p<0.001, one-way ANOVA Tukey test.

### Metabolomics revealed alterations in glucose-and lipid metabolites by STP treatment in U87 cells

STP’s role in metabolic inhibition in GBM is not clearly known despite suggestions that it is an LDH inhibitor. Therefore, we performed metabolomics analysis to identify the metabolic shifts and vulnerabilities of GBM cells after STP treatment. The volcano plot shows the top significantly altered metabolites in U87 cells after 24 h of treatment with STP (Fig. 2A). The downregulated metabolites included lithocholate, quinoline, succinyl coenzyme A, linoleic acid, uridine 5’ diphosphoglucose, erucic acid, 3-hrdroxyphenyl acetate/2,4 dihydroxy acetophenone whereas a higher expression of 3-methylhistamine/1 methylhistamine and trisodium 2-methylcitrate racemic mixture was observed (Fig. 2B). The top 25 metabolites besides the ones mentioned above included metabolites involved in lipid metabolism such as decreased carnitine which assists in turning fat to energy, erucic acid also involved in lipid metabolism and some interesting metabolites such as decrease in L-cysteic acid and cysteine sulfinic acid which can promote cancer cells to efficiently remove reactive oxygen species (ROS) and lead to cell cycle arrest. Catabolism of cysteine can also increase ROS production due to the regulation of glutathione. Several other decreased metabolites, such as Uridine-5-diphosphoglucose, thiamine monophosphate, etc., are responsible for the biosynthesis of nucleotide bases. Fig. 2C shows the dot plot of the significantly impacted metabolites, which includes linolic acid metabolism, vitamin B6 metabolism, glycerolipid metabolism, tryptophan metabolism, pyrimidine metabolism, purine metabolism, one-carbon pool by folate mechanism, and thiamine metabolism. These findings suggest a significant impact on lipid metabolism and an impact on DNA building blocks for cellular proliferation.

**Figure 2.**
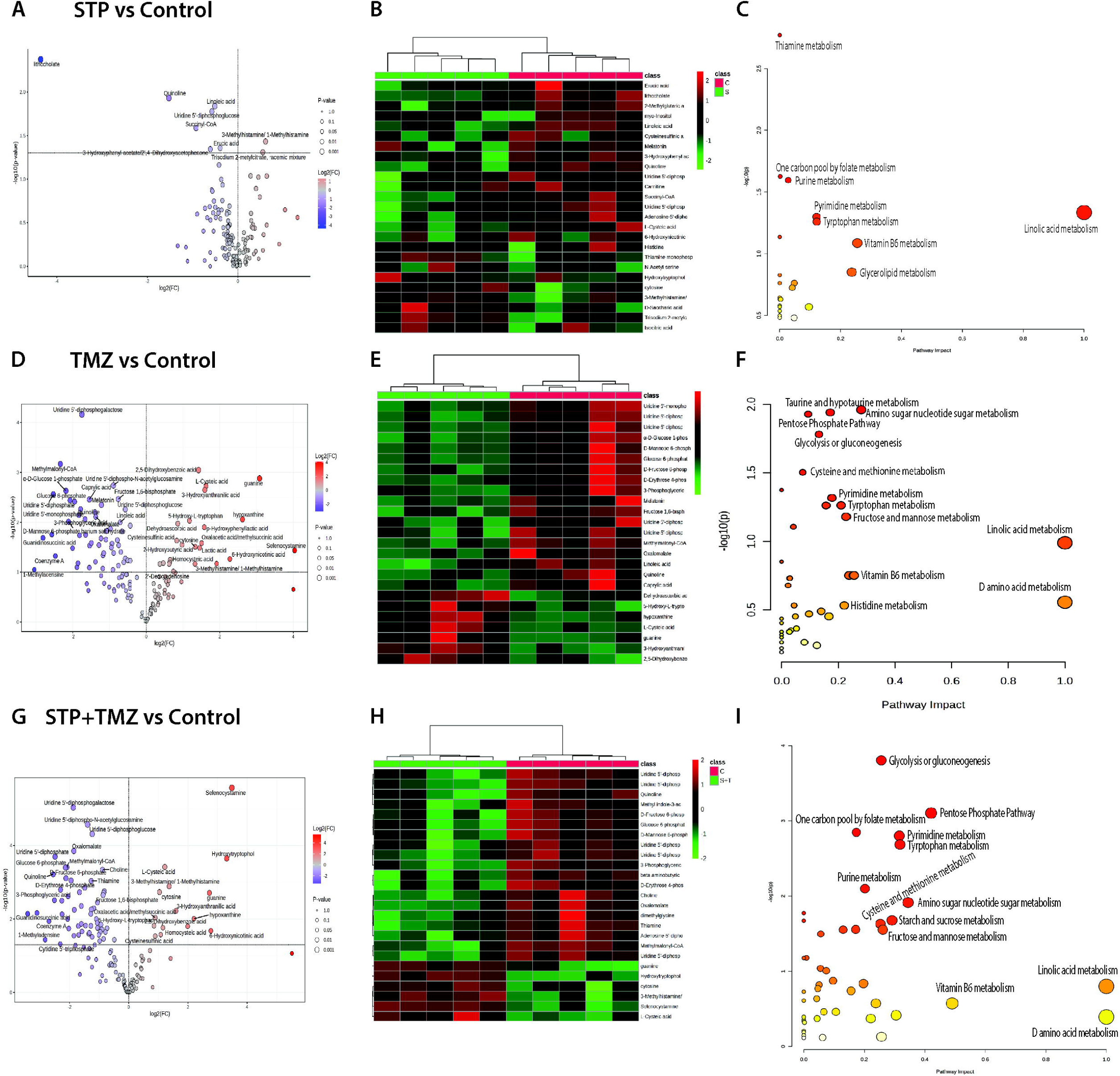
Metabolomics analysis reveals the altered metabolism in U87 cells. A), D) and G) volcano plots of STP vs Control, TMZ vs Control and STP+TMZ vs Control, depicting the major metabolites affected with treatment groups, respectively. B), E) and H) Heatmap analysis of top 25 metabolites affected in STP vs Control, TMZ vs Control and STP+TMZ vs Control groups, respectively. C), F) and I) Dot plot of Top 25 metabolic pathways involved in altered metabolism in treatment groups STP and combination of STP and TMZ, respectively. n=5, p<0.05.

### Combination treatment of STP and TMZ revealed global metabolic shifts in the biosynthesis of nucleotides, amino acids, sugars, and antioxidants

Interestingly, our combination treatment of TMZ and STP induced a drastic metabolic shift in the building blocks of DNA and RNA, including a decrease in uridine 5’ diphosphogalactose, UDP-glucose, thiamine, UDP, UMP, etc. Several metabolites in the combination treatment were strongly affected by TMZ treatment. We observed significant fold change differences in metabolites such as uridine 5’ diphosphogalactose, UDP-glucose, thiamine, UDP, UMP, etc. More changes in metabolic shift by TMZ treatment can be found in Fig. 2D. Moreover, several glycolytic and energy-producing metabolites were also affected significantly, including a decrease in glucose-6-phosphate, oxalomalate, ADP, 3-phosphoglyceric acid, alpha-D-glucose-1-phosphate, etc. Simultaneously, an increased expression of L cysteic acid, cysteinesulfinic acid, homocysteic acid, 3 methylhistamine, cytosine, selenocytosamine, hydroxytryptophol, hypoxanthine, hydroxyanthrallic acid etc. was observed as shown in the volcano plot and heat map in Fig. 2 G, and H. Fig. 2 I shows pathway impact scores and the log fold change in the metabolites involved in these pathways including amino sugar nucleotide sugar metabolism, glycolysis/gluconeogenesis, PPP, pyrimidine, and purine metabolism.

### Seahorse analysis confirms the alteration of glycolysis and mitochondrial metabolism by STP treatment

STP significantly impacted the glycolysis process in U87 cells after 24 h of treatment with 25 and 50 μM concentrations. Fig. 3A shows the significant reduction in the extracellular acidification rate of U87 cells upon STP treatment. The non-glycolytic acidification of starved U87 cells was unaffected by 25 μM of STP; however, 50 μM STP significantly reduced it. Upon addition of glucose, the glycolysis was significantly decreased in both STP 25 as well as 50 μM at 24 h. Moreover, upon addition of oligomycin, which is an ATP synthase inhibitor, shifting the ATP production to glycolysis, the glycolytic capacity of U87 cells at 25 and 50 μM was severely reduced.

**Figure 3.**
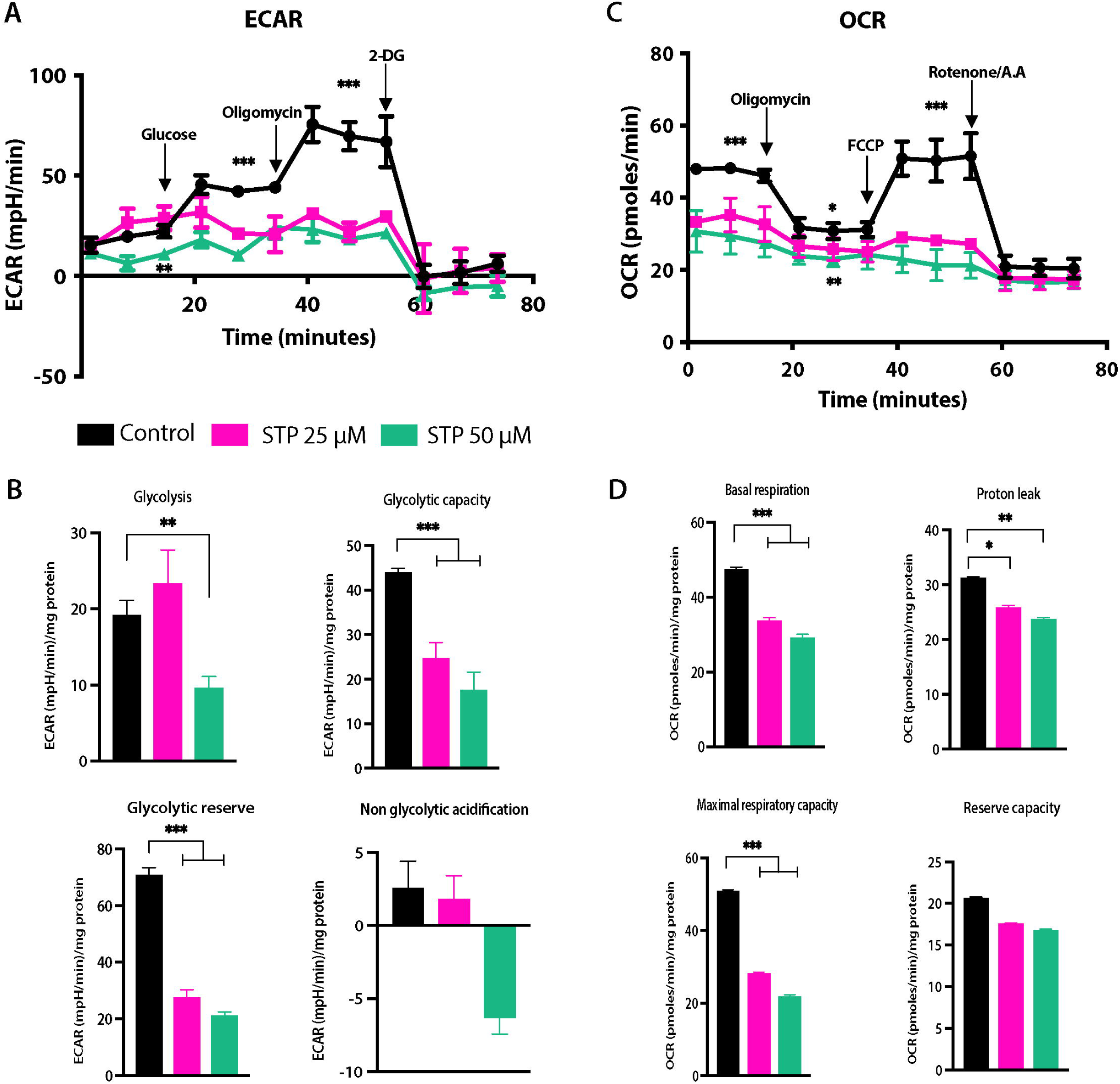
Seahorse metabolic analysis revealed the alterations in metabolism beyond glycolysis. A) Decreased Extracellular acidification rate (ECAR) on U87 cells upon 24 h treatment with STP at 25 and 50 mM concentration upon addition of glucose in port A, oligomycin in port B and 2 deoxy glucose in port C. B) Graphical representation of significant reduction in non-glycolytic acidification, glycolysis, glycolytic capacity and glycolytic reserve on STP treatment in U87 cells at 24 h. C) Decreased oxygen consumption rate (OCR) on U87 cells upon 24 h treatment with STP at 25 and 50 mM concentration after addition of oligomycin in port A, FCCP in port B, Rotenone and antimycin A in port C. D) Graphical representation of significant alteration in basic respiration, proton leak and maximal respiratory capacity in treatment groups STP 25 and 50 mM concentration. n=3, *p<0.033, **p<0.002, ***p<0.001, one way ANOVA Tukey test.

Thereafter, the addition of 2-deoxyglucose (2DG) caused a halt to glycolysis in the control and the STP treatment groups. Robust evidence of STP’s effect on glycolysis was thus observed. As we also had obtained many leads from metabolomics suggesting the involvement of the TCA cycle, we performed oxygen consumption rate (OCR) measurement. Our seahorse OCR assay results indeed validated the findings from metabolomics, suggesting that STP affects not only glycolysis but also OCR in GBM cells (Fig. 3C, D). The basal respiration of starved U87 cells was significantly affected, and a reduction was seen. The addition of oligomycin showed a significant proton leak and shifted cell metabolism towards glycolysis. Inhibition of complex I and III of the electron transport chain (ETC) by FCCP blocked the mitochondrial respiration of U87 cells altogether, and reserve capacity was unaffected in the STP treatment group.

### RNA seq unfolds key regulators of transcriptional changes in STP-treated and combination-treated GBM cells

Our metabolomic study uncovered that STP affected not only glycolysis but also cellular biosynthesis. Interestingly, some metabolites shed light on the auxiliary cell death mechanism, which included an increase in cysteine-like metabolites both in STP-and combination treatment. Therefore, we performed RNA seq to unravel the transcriptional changes in the STP-treated cells. Our results showed 74 significantly dysregulated genes expressed in STP-treated cells with an adjusted p-value of less than 0.05 (Fig. 4A). Some major downregulated genes included ALDH1A3, AOX1, SERPINB2, TXNIP, CPA4, SNAP25, etc. Some significantly upregulated genes included IL11 and KYNU (Fig. 4B). We also validated these findings by protein expression analysis of the differentially regulated genes by immunoblotting. As shown in Figure 6A, our findings corroborated with transcriptomic analysis, which strengthened our omics findings. Overall, we observed 1227 significantly differentially expressed genes in combination treatment in comparison to 685 significantly differentially expressed genes in TMZ-treated cells. Several genes affected by STP treatment were altered in STP+TMZ treatment, with higher fold changes, such as several stress-inducing genes ALDH1A3, AOX1, SERPINB2, CPA4. Whereas genes such as IL-1B, HSPA4L, MDM2, TRIM22 were some major genes common between TMZ and combination treatment (Figure 4D and G).

**Figure 4.**
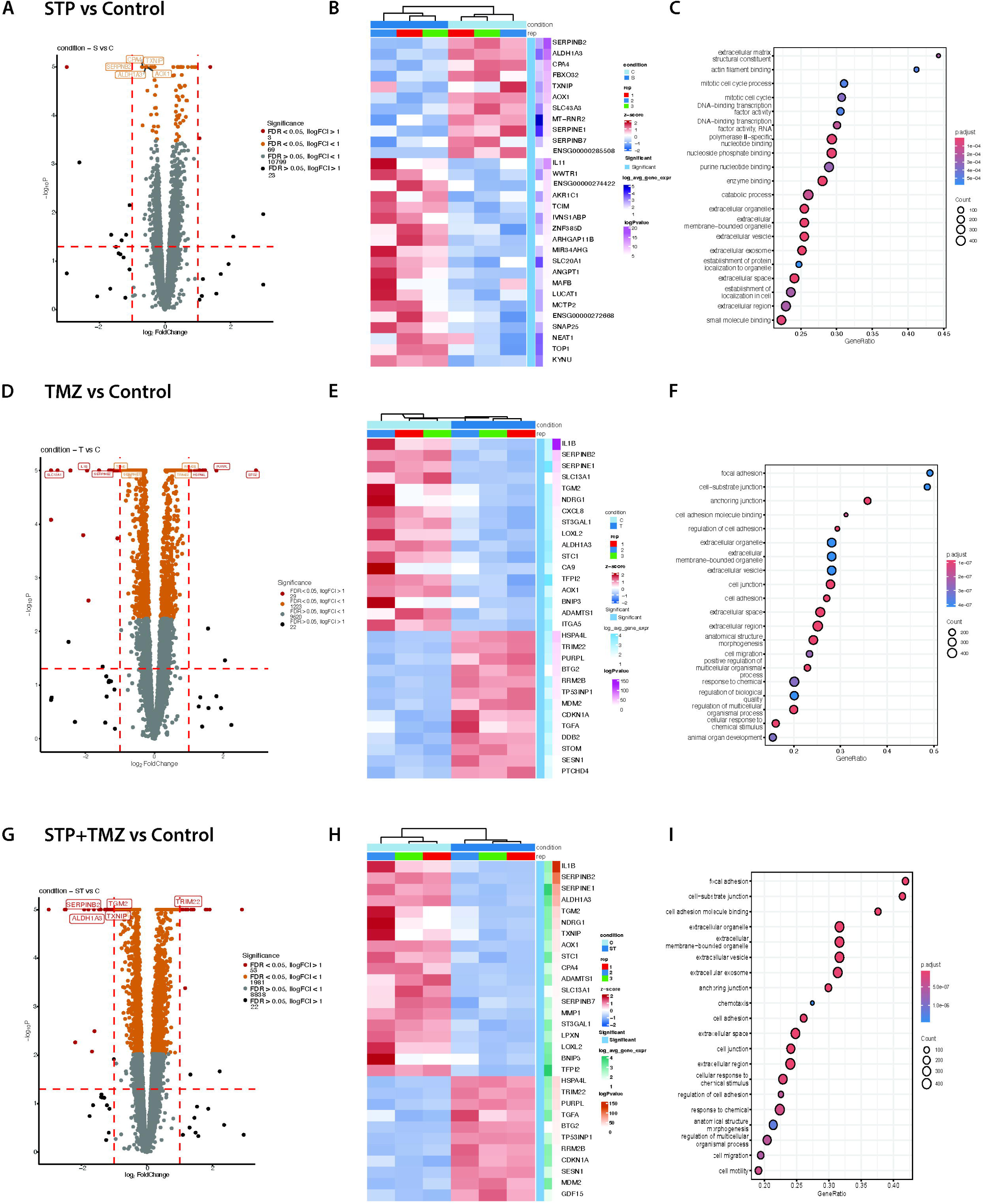
Transcriptomics analysis discloses the differentially expressed genes in U87 cells. A) D) and G) Volcano plot of STP vs Control, TMZ vs Control and STP+TMZ vs Control depicting the significantly differentially expressed genes affected with treatment groups, respectively. B), E) and H) Heatmap analysis of the top 30 genes affected in the STP vs Control, TMZ vs Control, and the STP+TMZ vs Control groups, respectively. C) F) and I) showing GSEA Gene ontology function highly affected with altered genes in STP+TMZ treatment. n=3, adjusted p<0.05.

### Metascape pathway analysis shows the major pathways involved in STP and the combination treatment effect on GBM cells

Metascape analysis results showed major involvement of nuclear events (kinase and transcriptomic factor activation), aldehyde metabolic process, pyridine nucleoside metabolic process, cellular catabolic process, metabolism of vitamins and cofactors such as NAD and NADP, and NOD-like receptor signaling pathway (Fig. 5A). The main GSEA pathway analysis showed us the involvement of genes in the mitotic cell cycle, cellular response to stress, and metabolism of carbohydrates. Remarkably, combination treatment showed interesting pathways involving VEGF and VEGFR2 signaling, PID p53 pathway, vasculature development, response to wounding, signaling to interleukins, positive regulation of programmed cell death, etc. All of which worked and contributed towards the cell death caused by the STP and TMZ combination treatment (Fig. 5C). TMZ treatment had a major role to play in the involvement of VEGF and VEGFR2 signaling, p53 pathway, as well as response to wounding and programmed cell death, which was noted in our control vs TMZ-treated analysis (Fig. 5B). GSEA pathway analysis also showed the downregulation of most genes involved in the VEGF pathway, which could lead to decreased angiogenesis. Moreover, several cancer pathways were involved, including p53, HIF1, and PI3K-Akt. The results also suggested that the immune response was impacted by the combination treatment, which may have also been a response to wound healing, as well as macrophage-stimulating protein MSP signaling.

**Figure 5.**
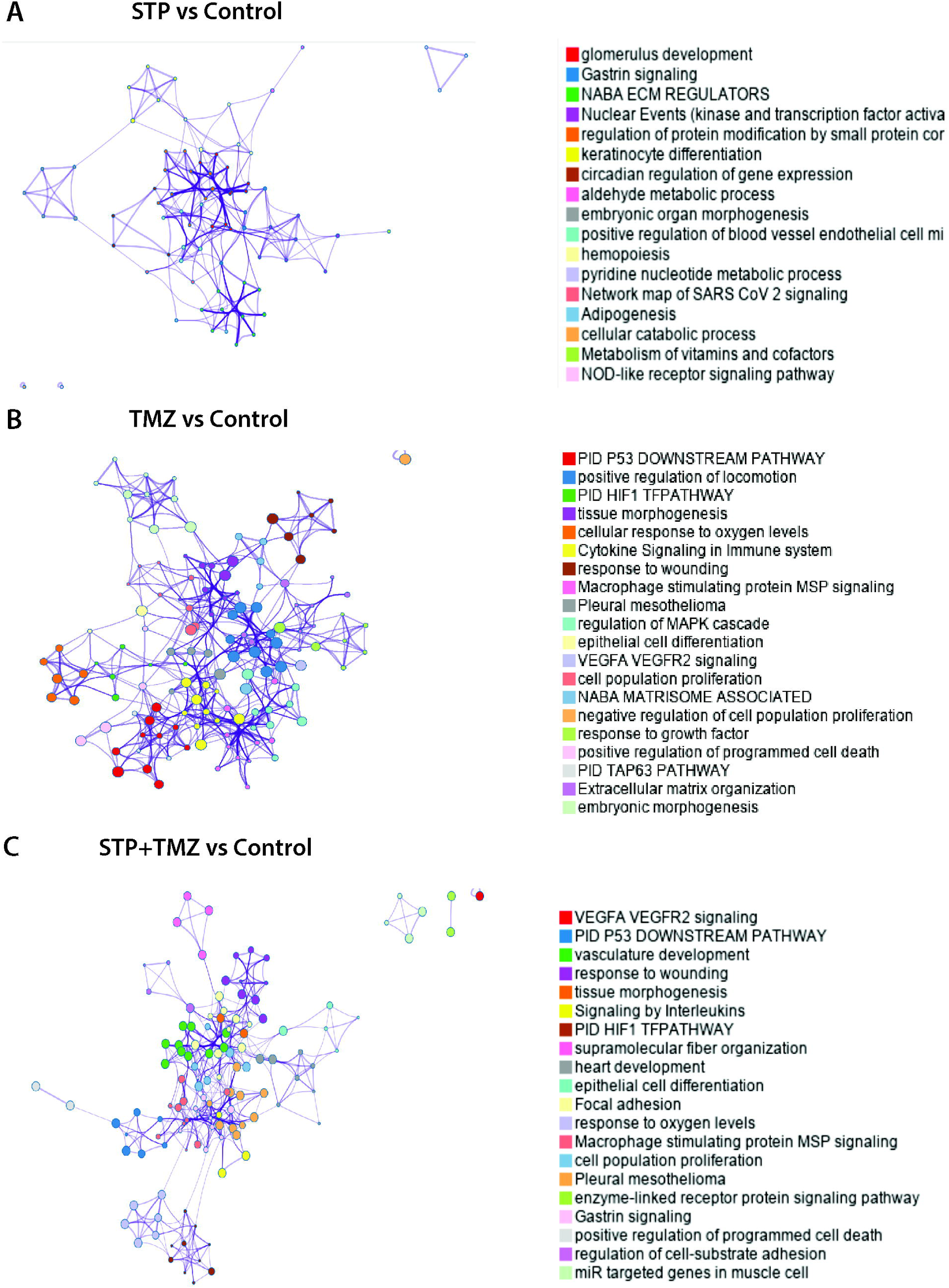
Transcriptomics analysis shows major pathways involved in differentially expressed genes in U87 cells. A) B) and C) Metascape pathway analysis of STP vs Control TMZ vs Control and STP+TMZ vs Control depicting the significantly altered pathways affected with treatment groups, respectively. n=3, adjusted p<0.05.

### RNA seq gene alterations were validated by protein expression analysis

The top significantly altered genes were then analyzed with protein expression analysis using immunoblotting. Decreased protein expression of ALDH1A3, AOX1, SERPINB2, and an increase in KYNU expression were observed, which corroborated our RNA seq findings. Significant decrease in CPA4 was observed in TMZ and STP+TMZ; however, the decrease in STP was moderate but not significant. An increased expression of TXNIP was observed across all three treatments STP, TMZ, and STP+TMZ; however, it was not significant (Fig. 6A and B).

**Figure 6.**
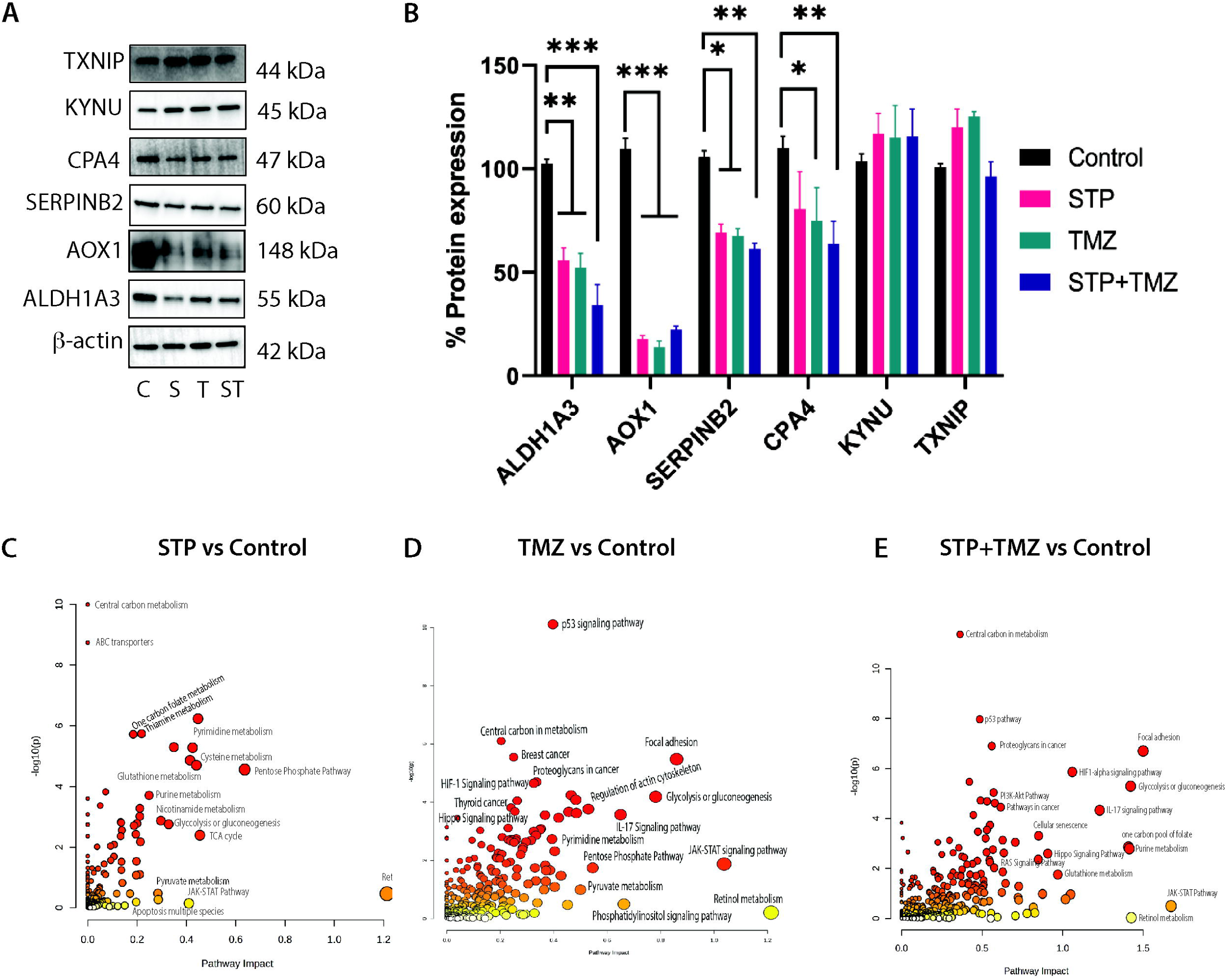
Validation of RNA seq analysis of top DEGs by protein expression analysis and Joint analysis showing major pathways involved in differentially expressed genes as well as altered metabolites in U87 cells. A) and B) Immunoblotting and quantification of protein expression, one-way ANOVA, ***p<0.001. C), D) and E) Dot plot analysis of STP vs Control, TMZ vs Control, and STP+TMZ vs Control, depicting the significantly altered pathways affected with treatment groups, respectively. n=5 metabolite list and n=3 gene list, p<0.05.

### Joint analysis of metabolomics and transcriptomics unravels the crosstalk between genes and metabolites

We performed a joint analysis of the differentially expressed genes and metabolites to evaluate the crosstalk between them and the pathways involved. Joint analysis of STP unfolded the combined outcome of transcriptional and metabolic alterations caused by the treatment on metabolic pathways. Fig. 6C shows the pathways affected, including retinol metabolism, tryptophan metabolism, nicotinate and nicotinamide metabolism, vitamin B6 metabolism, complement and coagulation cascades, hippo signaling pathway, NOD-like receptor signaling pathway, HIF1-alpha signaling pathway, degradation of amino acids like valine, leucine, and isoleucine, as well as the JAK-STAT pathway. Moreover, the combination of STP and TMZ treatment resulted in drastic alterations in the metabolic and transcriptional processes, including the p53 signaling pathway, focal adhesion, glycolysis, JAK-STAT signaling pathway, PPP, regulation of actin cytoskeleton, central carbon metabolism, and hippo signaling pathway. Although a lot of these pathways were predominantly affected by TMZ treatment (Fig. 6D and E), pathways such as Hippo signaling and the impact of proteoglycans in cancer stood out in the combination treatment. It would be interesting to study these mechanisms for STP’s impact on combination treatment with TMZ. Especially hippo signaling pathway has been highly associated with regulating organ size, cellular proliferation, and apoptosis.

### ROS levels were increased in STP-treated U87 cells

We performed a CMH_2_DCFDA staining assay on U87 cells to evaluate the ROS-associated stress by STP and combination treatment. We observed higher fluorescence in STP, combination groups, and positive control etoposide, indicating increased ROS levels (Fig. 7). The merged images show that approximately 90% of cells took up the CMH_2_DCFDA as compared to the control. These findings, therefore, validate that the mechanism of cell death in STP’s anticancer activity in GBM cells is associated with its ability to induce ROS.

**Figure 7.**
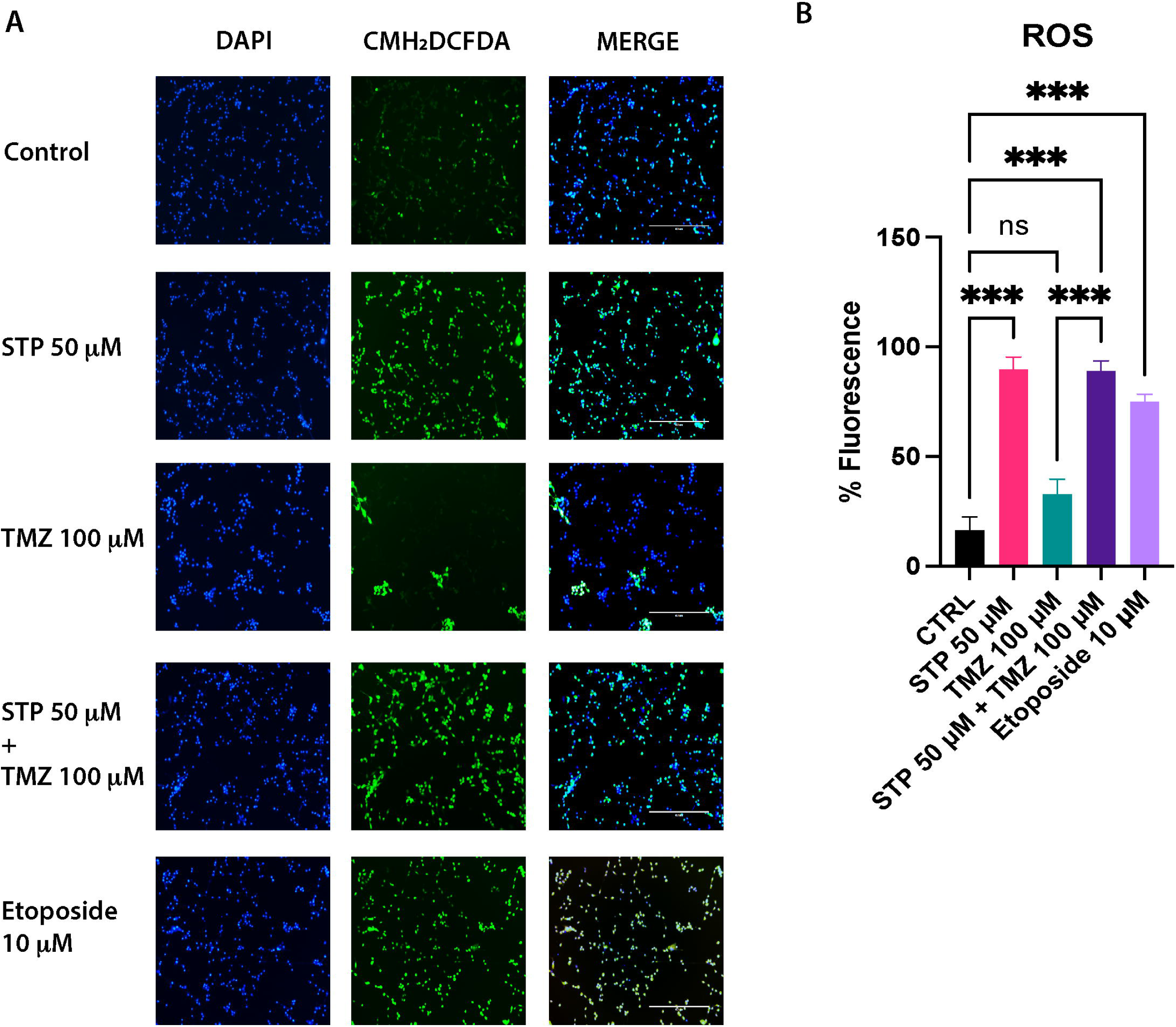
Increased ROS levels in treatment with STP and STP+TMZ treatment in U87 cells. A) Representative fluorescent images of cells treated with STP, TMZ, combination and positive control etoposide are shown in DAPI, CMH_2_DCFDA, and merge Magnification 100×, scale bar 100 mm. B) Quantification of the cells with CMH_2_DCFDA uptake with respect to DAPI from Image J and plotted on GraphPad Prism software, one-way ANOVA n=3, ***p<0.001.

### STP treatment resulted in decreased mitochondrial membrane potential (MMP)

CMXROS MMP assay was performed on U87 cells to evaluate the presence of functional mitochondria after STP treatment. In Fig. 8, decreased RFP fluorescence can be seen in STP, TMZ +STP combination-treated cells, and FCCP positive control compared to the control cells. The merged images show the DAPI-stained live cells with RFP-stained cells with regular MMP.

**Figure 8.**
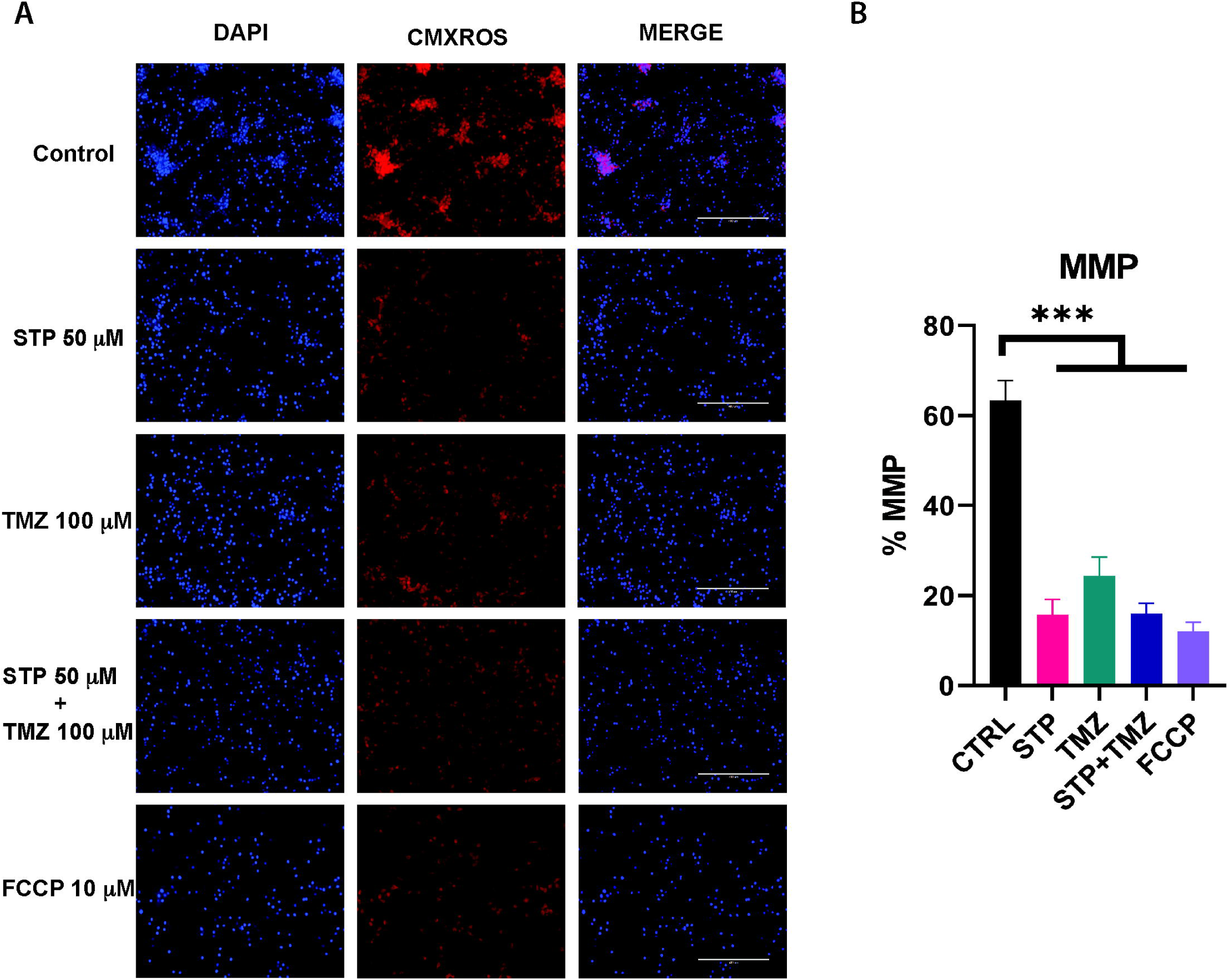
Decreased mitochondrial membrane potential with STP and STP+TMZ combination treatment in U87 cells. A) Representative fluorescent images of cells treated with STP, TMZ, combination, and positive control FCCP are shown in DAPI, CMXROS, and merge. Magnification 100×, scale bar 100 mm. B) Image analysis of fluorescent cells using Image J software and analysis using GrpahPad Prism, one-way ANOVA n=3, ***p<0.001.

### Phospho-Kinase array did not reveal any changes in the expression of 32 kinases

The phosphokinase array did not yield any significant alterations in the 32 kinases that are probed with STP lysates. No significant changes in phosphorylation profiles of proteins that are conventionally relevant in cancer-associated pathways, such as EGFR, ENOS, ERK1/2/3, FGR, JNK, AKT1/2/3, and C-JUN, were observed Fig. 9A and B. This ruled out several mechanisms that could potentially be behind STP’s anti-cancer activity. We also evaluated autophagy and lipid peroxidation-associated ferroptosis by STP treatment, and our results did not show autophagy or ferroptosis-associated cell death, as shown in Fig. 9 C, D and E.

**Figure 9.**
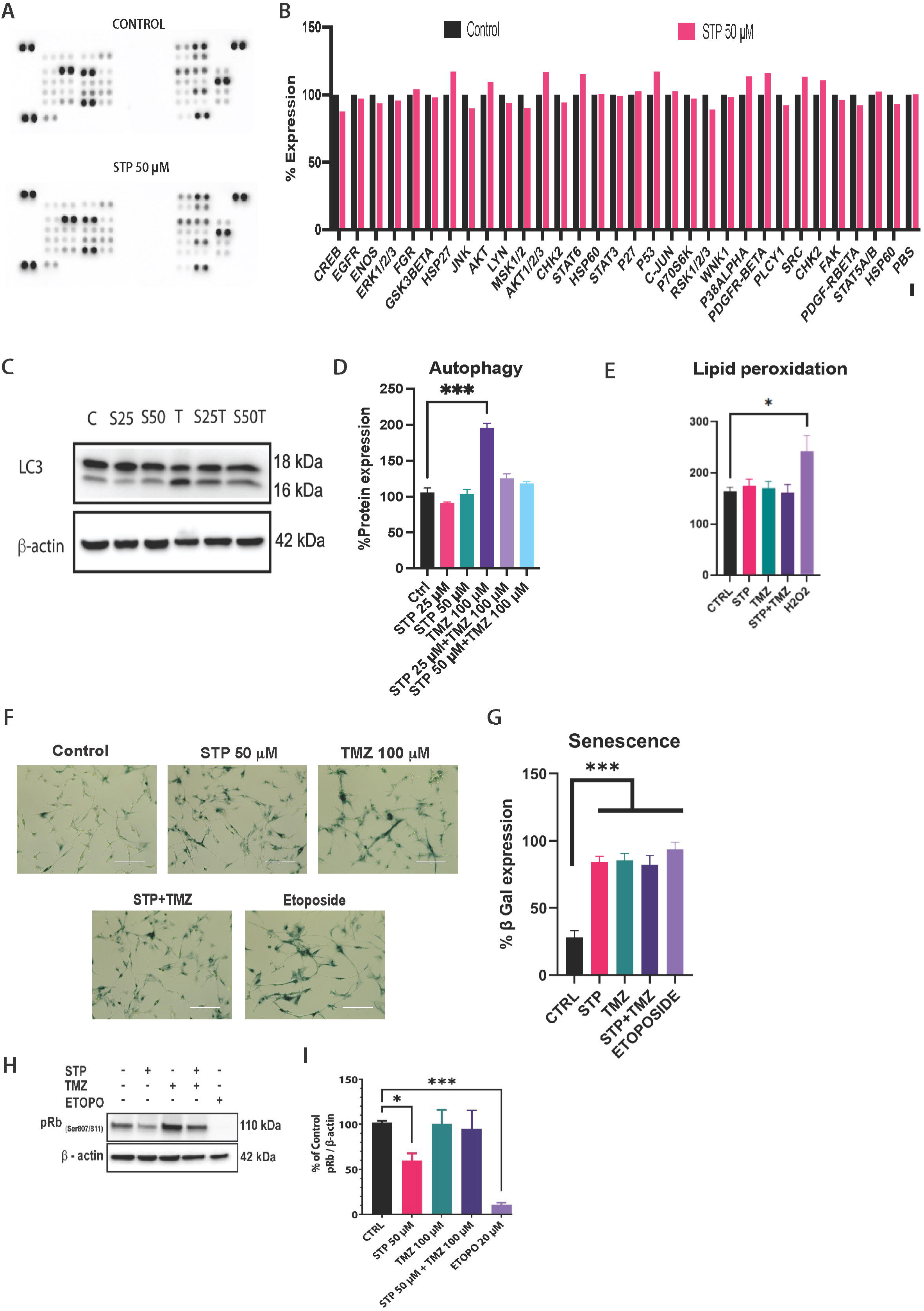
Increased senescence but no significant change in major kinases, autophagy, and lipid peroxidation in treatment with STP in U87 cells. A) Kinase array showing 32 major kinases and B) its quantification. C and D) Immunoblot of autophagy marker LC3 and its quantification. E) Quantification of MDA by lipid peroxidation. F) Representative colorimetric images of β-galactosidase uptake in U87 cells after 48 h treatment with STP, TMZ, combination of STP+TMZ, and etoposide positive control. Magnification 200×. Scale bar 200 mm. G) Quantification of senescent cells using Image J and GraphPad Prism software, one-way ANOVA, n=3, ***p<0.001. H) Representative western blots depicting phospho-Rb in U87 cells after 48 h treatment with STP, TMZ, combination, and etoposide as a positive control. I) Quantification of pRb blot using Image J and analysis using GraphPad Prism software, one-way ANOVA, n=3, *p<0.033, ***p<0.001.

### STP treatment induces cellular senescence in U87 cells

STP and its combination with TMZ were evaluated for their ability to induce senescence in U87 cells. Senescence, which is characterized by irreversible cell cycle arrest and higher expression of β-galactosidase (SA-β-gal), was shown to increase in STP-treated cells as well as the STP+TMZ combination (Fig. 9 E). In addition, a decreased expression of phospho-Rb was observed in STP group alone whereas in combination group there was no significant difference (Fig. 9 H and I). These results, combined with ROS levels as well as our prior findings on cellular proliferation and lack of robust apoptosis, support that STP-mediated death of U87 cells involves senescence [10]. In addition, ROS-mediated autophagy was studied by studying LC3 protein expression, and we did not observe significant alterations by STP treatment (Fig. 9 C and D).

## Discussion

In this study, we identified key mechanistic effects of STP-induced cytotoxicity on GBM cells. We employed a multiomic approach studying metabolomics and transcriptomics coupled with Seahorse metabolic studies and cell biology assays to mechanistically study STP’s anticancer effects in U87 cells. Our study demonstrated that STP treatment results in metabolic alterations of both glycolysis and the TCA cycle, resulting in decreased MMP, increased ROS levels, and senescence-mediated cell death.

STP, an FDA-approved anti-epileptic drug for Dravet syndrome, has lately been studied for its role as a putative LDH inhibitor [11,12]. A study conducted across 80 NSCLC cell lines analyzed the nutrient consumption and addiction-promoting cell growth. In their study, they observed higher accumulation and uptake of lactate during glycolytic activity that proved that lactate has a multifunctional signaling role and is not just a waste byproduct. Another study on tumor microenvironment (TME) for cancer progression recognized that in response to hypoxic conditions, cells generated a large amount of lactate by metabolizing glucose and glutamine. This lactate, coupled with the release of H^+,^ prevents the intracellular acidification but promotes TME acidification. Lactate is reported as a tumor-promoting metabolite that metabolically reprograms cancer cells. LDH enzyme catalyzes the reversible conversion of pyruvate to lactate and plays a pivotal role in cancer metabolism [15]. LDH has been studied extensively for its upregulation in highly progressive cancers with low survival and poor prognosis [16].

Tumor cells upregulate glycolysis even in the presence of oxygen. This phenomenon, known as the Warburg effect or aerobic glycolysis, is an established hallmark of cancer. Although it leads to the generation of fewer ATPs as compared to the TCA cycle, it yields several cofactors, such as NAD+ and NADH, that can be utilized in various biological processes and to synthesize building blocks of amino acids and nucleic acids. In our study, we not only explored the role of STP as an LDH inhibitor but also dove deep into the mechanistic targets of STP’s anticancer activity in GBM. Our studies revealed that although STP followed similar trends to other known LDH inhibitors, such as GSK2837808A and Na Oxamate in LDH overexpressed and silenced cells, it proved to be a moderate LDH inhibitor in our enzyme activity assay (Fig. 1). Our previous study has shown the synergy between STP and TMZ as a combination for the treatment of GBM [10,17]. In this study, we mechanistically investigated both STP and the STP+TMZ combination in GBM cells using a multiomic approach. The Seahorse assays revealed that STP strongly decreased the ECAR and OCR levels in U87 cells, indicating that not only glycolysis but also oxidative phosphorylation was significantly affected (Fig. 3C). The mitochondrial membrane potential results are a validation of the dysregulated functioning of mitochondria in STP-treated cells (Fig. 8).

Our metabolomic analysis revealed the involvement of STP in various pathways, including linoleic acid metabolism, pyrimidine metabolism, tryptophan biosynthesis, and the one-carbon pool by folate. Some of these metabolites, such as lithocholate, linoleic acid, and succinyl coenzyme A, appear intriguing due to their implied importance in GBM. Higher expression of lithocholate is often reported to cause apoptosis in cancer cells such as neuroblastoma and glioblastoma [18]. An increase in linoleic acid suggested lipid metabolism involvement in the process. Although the role of linoleic acid in GBM is not well established, some studies show that the presence of excess linoleic acid can alter tumor cell proliferation, migration, and apoptosis [19]. These findings indicate the compensatory mechanism of U87 cells to resist cell stress or increased lipid remodeling or an adaptive mechanism to resist oxidative stress. Succinyl coenzyme A-a major metabolite involved in the process of TCA cycle suggests that STP may not only affect glycolysis but also oxidative phosphorylation. The lack of succinyl co enzyme A can lead to interference in important processes of succinylation in glucose metabolism, amino acid metabolism, fatty acid metabolism, ketone body synthesis, and reactive oxygen species clearance [20]. It is known that succinylation can promote tumorigenesis and the posttranslational modification (PTM) leading to tumor cell proliferation and development in gliomas requiring succinyl co enzyme A [20].

To evaluate the role of genetic vulnerabilities of GBM along with metabolic reprogramming, we studied the transcriptomic landscape of U87 cells upon treatment with STP, TMZ, and their combination. Although STP did not cause major transcriptional differences, it led us to some clues for its additive action besides the metabolic inhibition of LDH and affecting energy generation pathways. Our major hits included ALDH1A3, AOX1, TXNIP, CPA4, SERPINB2, FGF1 and KYNU to name a few. ALDH1A3 and AOX1 have recently been studied for their role in ROS-associated cell death in cancer cells [21]. Decrease in ALDH1A3, also known as aldehyde dehydrogenase 3, renders cells more vulnerable to glycolysis inhibition and affects cell migration [21]. Our earlier studies have reported the effect of STP on decreased migration in GBM cells. Overexpression of AOX1 has previously been reported to promote proliferation, migration, and invasion of gallbladder cancer [22]. AOX1 overexpression also promoted cell proliferation and invasion in colorectal cancer and inhibited apoptosis [23]. TXNIP plays a pivotal role in the redox reactions. The role of TXNIP is not clearly understood in cancer, as some studies suggest it acts as a tumor suppressor for cancers such as breast cancer, bladder cancer, and gastric cancer [24]. Whereas another study showed that higher expression of TXNIP was associated with angiogenesis and relapse in renal cell carcinoma [25]. TXNIP’s role in cancer remains complex; however, it is certain that it impacts ROS production. The CPA4 gene, which also acts as a biomarker for cancer progression, survival, and growth, is associated with cancers such as gastric cancer, pancreatic cancer, etc, and its knockdown caused increased survival or decreased tumor growth [26,27]. These genes in the combination treatment group with TMZ were highly significantly affected compared to STP alone. This indicated that STP may boost TMZ’s anti-cancer effect, and an increase in ROS levels can be one of the major mechanisms that led to STP’s anti-cancer activity, as suggested by metabolomics and transcriptomics. We validated our hits using immunoblot protein expression. Our joint analysis showed disruptions in retinol metabolism. Retinol metabolism has been increasingly recognized for its role in cancer cell differentiation, proliferation, and oxidative stress regulation. ALDH1A3 is a major enzyme responsible for oxidation of retinaldehyde to retinoic acid and retinol metabolism pathway. This process also generates retinoic acid which regulates gene expression associated with cell proliferation, differentiation and survival. Moreover, it contributes to detoxification of reactive oxygen species (ROS).

Interestingly, our hits leaned us towards ROS-induced cellular stress as STP’s contribution to GBM cell death. We, therefore, performed ROS fluorescent staining using CH_2_DCFDA and DAPI. STP increased the ROS levels in U87 cells. Given that metabolic inhibitors have been shown to impact cell fate decisions by modulating differentiation-associated pathways [28], it is plausible that STP exerts additional effects beyond direct metabolic disruption. A decrease in cysteine triggers ferroptosis-mediated cell death by degradation of GPX4 [29]. Therefore, we aimed to study the cell death mechanism of STP treatment in GBM cells. For this, we systematically investigated multiple well-characterized cell death pathways by phospho kinase array and explored pathways such as apoptosis, ferroptosis, and autophagy. Our analysis revealed negative results for all three. Our previous findings have demonstrated that STP did not induce apoptosis in U87 cells [10], and we observed that it did not affect major cancer-associated kinases (Fig. 9 A and B). We also studied the effect of STP on lipid peroxidation-associated ferroptosis as well as autophagy. Our results indicated that STP does not cause lipid peroxidation-associated ferroptosis and showed no changes in autophagy marker LC3 (Fig. 9 C and D). In contrast, we observed a robust induction of cellular senescence, as evidenced by an increase in β-galactosidase activity and a significant decrease in phospho-Rb levels (Fig. 9 I). Disruption of retinol metabolism is a known mechanism to increase ROS production. Elevated ROS triggers cellular senescence, primarily through induction of DNA damage response, which also confirms DNA damage response-associated significant transcriptional alterations in biological processes such as DNA binding transcriptional activity, nucleoside binding, etc. Similarly, the downregulation of metabolites UMP, UTP, UDP-glucose, and thiamine strongly suggests the process of cellular proliferation has halted, which again can be validated by the effect on the mitotic cell cycle through transcriptional changes.

Although our study successfully demonstrated STP’s mechanism of cellular senescence and inhibition of cellular proliferation, there are some limitations. The precise molecular mechanisms and signaling involved through which STP exerts its activity remains unclear. We have performed efficacy on multiple cell lines previously; however, our mechanistic analysis was performed on U87 cells, and it would be interesting to evaluate the same in other cell lines to increase applicability. Collectively, our findings suggest that GBM cells after STP treatment, rather than undergoing conventional cell death, predominantly enter a state of senescence driven by metabolic reprogramming, oxidative stress, and mitochondrial dysfunction. This study highlights the complex interplay between metabolic and transcriptional pathways.

## Acknowledgements

This study was supported by an award from the National Institute of General Medical Sciences of the National Institutes of Health under Award Number 5R16GM145557 to Vikas V. Dukhande and by funds from the College of Pharmacy and Health Sciences, St. John’s University, Queens, NY, USA. Authors would also like to thank the Einstein-Mount Sinai Diabetes Research Center’s isotope and metabolomics core facility and Yale Center for Genome Analysis for help with metabolomic and transcriptomic studies, respectively.

## CRediT authorship contribution statement

Yadav Anjali: Writing – original draft, Formal analysis, Data curation, Conceptualization. Bhutkar Shraddha: Data curation, Formal analysis. Barot Shrikant: Data curation, Formal analysis. Dukhande Vikas: Resources, Funding acquisition, Conceptualization, Review and editing, Supervision, Project administration.

